# High-throughput single nucleotide polymorphism (SNP) discovery and validation through whole-genome resequencing of hundreds of individuals in Nile tilapia (*Oreochromis niloticus*)

**DOI:** 10.1101/594671

**Authors:** J.M. Yáñez, G. Yoshida, A. Barria, R. Palma-Véjares, D. Travisany, D. Díaz, G. Cáceres, M.I. Cádiz, M.E. López, J.P. Lhorente, A. Jedlicki, J. Soto, D. Salas, A. Maass

## Abstract

Nile Tilapia (*Oreochromis niloticus*) is the second most important farmed fish in the world and a sustainable source of protein for human consumption. Several genetic improvement programs have been established for this species in the world and so far, they are mainly based on conventional selection using genealogical and phenotypic information to estimate the genetic merit of breeders and make selection decisions. Genome-wide information can be exploited to efficiently incorporate traits that are difficult to measure in the breeding goal. Thus, SNPs are required to investigate phenotype–genotype associations and determine the genomic basis of economically important traits. We performed *de novo* SNP discovery in three different populations of farmed tilapias. A total of 29.9 million non-redundant SNPs were identified through Illumina (HiSeq 2500) whole-genome resequencing of 326 individual samples. After applying several filtering steps including removing SNP based on genotype and site quality, presence of Mendelian errors, and non unique position in the genome, a total of high quality 50,000 SNP were selected for validation purposes. These SNPs were highly informative in the three populations analyzed showing between 43,869 (94%) and 46,139 (99%) SNP in HWE; 37,843 (76%) and 45,171(90%) SNP with a MAF higher than 0.05 and; 43,450 (87%) and 46,570 (93%) SNPs with a MAF higher than 0.01. The final list of 50K SNPs will be very useful for the dissection of economically relevant traits, enhancing breeding programs through genomic selection as well as supporting genetic studies in farmed populations Nile tilapia using dense genome-wide information.

## INTRODUCTION

The study of phenotype-genotype association, identification of the genomic basis of economically important traits and the implementation of genomic predictions in farmed fish require a considerable number of highly informative single nucleotide polymorphisms (SNP) that preferably segregate in multiple populations. Thus, the discovery and characterization of dense SNP panels will help to a better understanding of complex traits architecture and genome biology in farmed fish (Yáñez et al. 2015). From an animal breeding perspective, the use of a high number of SNPs markers to support tilapia genetic improvement programs has the potential to speed up genetic gains for traits which by their nature cannot be directly recorded in selection candidates e.g. carcass quality and disease resistance traits (Yáñez and Martinez 2010; Ødegård et al. 2014; Yáñez et al. 2014). Dense SNP panels can also allow the determination of genomic regions underlying selection and adaptation to different environmental conditions during the domestication process in farmed fish populations (López et al. 2015).

The discovery of SNP markers in aquaculture species of commercial interest has been widely spread due the availability of high quality reference genomes as it is the case of Atlantic salmon (*Salmo salar*) (Lien et al. 2016), rainbow trout (*Oncorhynchus mykiss*) (Berthelot et al. 2014) and Pacific oyster (*Crassostrea gigas*) (Zhang et al. 2012). These information has facilitated the development of dense SNP panels, being currently available for different species including Atlantic salmon (Houston et al. 2014; Yáñez et al. 2016), rainbow trout (Palti et al. 2015), channel catfish (*Ictalurus punctatus*) (Liu et al. 2014; Zeng et al. 2017) and Pacific oyster (Gutierrez et al. 2017; Qi et al. 2017). These genomic resources have been used to carry out several studies aimed at identifying the genetic architecture of economically relevant traits in fish by means of genome-wide association studies for traits such as growth (Gutierrez et al. 2015; Tsai et al. 2015; Yoshida et al. 2017; Neto et al. 2019), disease resistance (Correa et al. 2015; Tsai et al. 2016; Barría et al. 2018) and carcass quality (Gonzalez-Pena et al. 2016). These SNP panels have also been used to test different approaches for the implementation of genomic predictions in Atlantic salmon (Ødegård et al. 2014; Bangera et al. 2017; Correa et al. 2017; Sae-Lim et al. 2017) and rainbow trout (Vallejo *et al.* 2016, 2017, 2018; Yoshida, *et al.* 2018a; Yoshida, *et al.* 2018b).

Nile tilapia (*Oreochromis niloticus*) is among the most important fresh-water species farmed worldwide. Several selective breeding programs have been stablished for this specie since 1990’s, allowing to genetically improve important commercial traits and expand tilapia farming across the globe. To date, the most widespread improved tilapia strain is the Genetically Improved Farmed Tilapia (GIFT) (Webster and Lim 2006), being farmed in Latin America, Asia, and Africa (Gupta and Acosta 2004). It has been shown that the response to selection for growth rate reached up to a 15% per generation after six generations of selection (Ponzoni et al. 2011), demonstrating the feasibility to improve this trait by means of artificial selection. However, and despite the large number of genetic programs and the advantages of Nile tilapia farming (e.g. fast growth and high adaptability), there are scarce studies on the application of genomic technologies for mapping variants associated with desired traits and enhancing selection through the use of genomic predictions in comparison with other aquaculture species. Consequently, up to date, genetic improvement programs mainly rely on traditional pedigree-based breeding approaches, with only one published report on the development of genome resources to enhance selective breeding in a single Nile tilapia population (Joshi et al. 2018).

The objective of this study was to perform a large-scale *de novo* SNP discovery and using whole genome resequencing of hundreds of Nile tilapia individuals from three different farmed populations and characterize a medium-density (50K) SNP panel to be further used in the determination of the genetic basis of complex traits and genomic selection in this species.

## MATERIALS AND METHODS

### Populations

The principal aim of the present study was to discover and characterize highly informative SNP variants in Nile tilapia farmed populations. Thus, we included animals from three different commercial breeding populations established in Latin America, originated from admixed stocks imported from Asia. We used 59 samples from POP A breeding population (Brazil) and 126 and 141 samples from POP B and POP C breeding populations, respectively, both belonging to Aquacorporación Internacional (Costa Rica). The three breeding populations are directly or indirectly related to the GIFT (Genetically Improved Farmed Tilapia), which is the most spread Nile tilapia strain used for farming purposes worldwide. The GIFT strain was initially established in Philippines by the crosses between four farmed Asian strains originally from Israel, Singapore, Taiwan and Thailand and four wild strains from Egypt, Senegal, Kenya and Ghana. The POP A breeding population represents GIFT animals which were introduced to Brazil for multiplication and farming purposes in early 2000. The POP B breeding population is a mixture of the original Asian farmed populations from Israel, Singapore, Taiwan and Thailand present in the Philippines in the late 1980s, which give origin to the GIFT strain. The POP C breeding population represents a combination of genetic material from the best available stocks corresponding to GIFT (Generation 8) and two original African strains founding GIFT. The three populations have been genetically improved for growth rate for more than 10 generations in total, using genetic evaluations based on the best linear unbiased predictor.

### Whole-genome resequencing

Tissue samples from the 326 fish were obtained by partial fin-clipping of fish anesthetized using benzocaine. Tissue sampling was carried out in accordance with the commercial practice and norms held by the two companies, Aquacorporación Internacional and Aquamerica, which provided the samples. Genomic DNA was extracted from fin-clip samples using the DNeasy Blood & Tissue Kit (QIAGEN) following manufacturer's protocol. Whole-genome resequencing was performed on each of the individuals multiplexing five bar-coded samples per lane of 150 bp paired-end in Illumina HiSeq-2500.

### SNP discovery and annotation

We used the assembly ASM185804v2 (GenBank accession GCF_001858045.1) of the *O. niloticus* as a reference genome sequence. This assembly consists of 1,010 Mb of total sequence comprising 2,990 contigs with a contig N50 of 3.09 kb. Sequences from all samples were evaluated using FASTQC (Wingett and Andrews 2018) to assess base quality and primer adapter contamination. Burrows-Wheeler Aligner (BWA-MEM) (Li and Durbin 2009) was used to map the reads of each sample to the reference genome. Briefly, BWA-MEM starts a local alignment between a fragment of the read to the reference genome and extends it until the read is completely mapped, if the read cannot be fully mapped is soft clipped or eventually discarded. To avoid invalid flags in further analysis, reads without a pair were discarded from the output using SAMtools (Li et al. 2009). In order to obtain a high quality BAM file all duplicated reads were masked as such using PICARD (http://broadinstitute.github.io/picard). For variant calling we used the standard protocol implemented in the Genome Analysis Tool (GATK) version 3.5.0. All high-quality BAM files for each sample obtained previously, were assessed at the SNP calling step and summarized into a single Genotyped Variant Calling Format file (VCF) containing all data. Each SNP was categorized as being either homozygous or heterozygous for the ALT allele (i.e., the non-REF allele). To call a sample homozygous for an ALT allele at a given site, the most common ALT allele variant confidence divided by the total sample reads (QD) must be at least 10 (QD > 10). This normalized ALT alleles in zones with high density depth and poor-quality calls. Only bi-allelic SNPs were pre-selected in posterior filters. The final VCF file was annotated using Variant Effect Predictor (VEP v92.1) in offline mode using the cached Orenil1.0 genome database and the gff file GCF_001858045.1_ASM185804v2_genomic.gff.

### SNP filtering and validation

Population genetics analyses and filtering described here, including Hardy-Weinberg Equilibrium (HWE), minor allele frequencies (MAF) and observed and expected heterozygosities (H_O_ and H_E_, respectively), were carried out using *VCFTools* (Danecek et al. 2011) and *Plink* (Purcell et al. 2007). An initial common quality control (QC) for the three populations was performed using VCF tools software. The genotypes were filtered to remove indels and sequence alterations, markers with genotypes quality (GQ<0.15) and minimum quality (minQ<40). Furtherly, specific QC filters for each population were applied discarding SNPs with missing genotypes >0.60, minor allele frequency (MAF) <0.01, Hardy-Weinberg Equilibrium (HWE) p-value < 1e-06, Illumina score <0.8 (Table 1). SNPs were also filtered based on Mendelian error using genotypes from 8 trios (sire, dam and offspring) from POP B, in which markers with less than one Mendelian error were retained. In addition, SNP probes were aligned to the Nile tilapia reference genome to retain markers which have a unique position in the genome assembly (GenBank accession GCF_001858045.1) generated by the University of Maryland and the University of Stirling (Conte et al. 2017) using the following procedure: i) SNP probes of 121 bp were built using flanking SNPs sequences (60 bp upstream and 60 bp downstream of each SNP); and ii) each probe was aligned to the reference genome by means of BLASTN version 2.3.0+ (Madden 2002), using the following parameters: word size of 11 (-w) and minimum e-value of e-40 (e); iii) all hits were evaluated tolerating only 2 mismatches, and no gaps were allowed; and iv) probes having a unique location in the genome were retained. All this procedure was achieved using in-house Python scripts. Furtherly, SNPs with MAF > 0.05 in the three commercial populations were prioritized. Finally, SNPs were selected so that they are as evenly distributed along the genome as possible. This was done by selecting SNPs from windows of equal size across various chromosomes of the genome using THIN < 9 kb command. When selecting SNPs from windows, higher preference was given to common SNPs between population POP B and POP C.

**Table 1.** Summary of results from SNP discovery and quality control filtering for SNP selection in 326 whole-genome sequenced individuals from three farmed Nile tilapia (*Oreochromis niloticus*) populations.

| Chr<sup>a</sup> | Initial<br>Number of<br>SNP | First<br>common<br>filter<sup>b</sup> | Population specific filters<sup>c</sup> |  |  | Last<br>common<br>Filter<sup>d</sup> |
| --- | --- | --- | --- | --- | --- | --- |
|  |  |  | POPA | POP B | POP C |  |
| Mito<sup>e</sup> | 1671 | 1596 | 0 | 0 | 0 | 0 |
| LG1 | 779881 | 708381 | 6768 | 31144 | 18046 | 2045 |
| LG2 | 891811 | 797928 | 7765 | 32143 | 12664 | 1881 |
| LG3a | 626179 | 539885 | 6583 | 17175 | 9991 | 714 |
| LG3b | 3491391 | 2943480 | 22758 | 47028 | 29881 | 1823 |
| LG4 | 1099494 | 983024 | 11401 | 38514 | 23132 | 2150 |
| LG5 | 792857 | 706930 | 6497 | 26718 | 11529 | 1811 |
| LG6 | 1247549 | 1112177 | 10300 | 47240 | 23576 | 2560 |
| LG7 | 1326771 | 1191205 | 16277 | 61818 | 24935 | 3549 |
| LG8 | 834050 | 752753 | 10587 | 35798 | 17751 | 1803 |
| LG9 | 842634 | 757126 | 7171 | 27404 | 13135 | 1518 |
| LG10 | 767756 | 689181 | 8389 | 29290 | 13115 | 1774 |
| LG11 | 904270 | 813207 | 8839 | 31506 | 16411 | 2074 |
| LG12 | 1178720 | 1051496 | 12065 | 39106 | 18552 | 2181 |
| LG13 | 797314 | 722353 | 7152 | 29682 | 18623 | 1819 |
| LG14 | 960008 | 861238 | 11750 | 39640 | 17386 | 2176 |
| LG15 | 991645 | 895658 | 10517 | 32442 | 19090 | 2002 |
| LG16 | 1319450 | 1181919 | 12268 | 49954 | 25188 | 2447 |
| LG17 | 1023476 | 919757 | 9226 | 40200 | 25573 | 2502 |
| LG18 | 1111651 | 988338 | 10630 | 39619 | 18910 | 2082 |
| LG19 | 624640 | 563099 | 7781 | 26643 | 15163 | 1799 |
| LG20 | 762408 | 685589 | 7586 | 27546 | 13409 | 2013 |
| LG22 | 1202188 | 1062702 | 13440 | 42665 | 18451 | 2016 |
| LG23 | 1229543 | 1099828 | 11769 | 43157 | 25489 | 2610 |
| **US<sup>f</sup>** | 5149044 | 4386247 | 24031 | 50640 | 31645 | 2651 |
| **Total** | **29956401** | **26415097** | **261550** | **887072** | **461645** | **50000** |

<sup>a</sup> Chromosome
<sup>b</sup> First common quality control filtering using all populations including SNP *excluded by genotypes* *quality* < 15 and minimum site *quality* < 40
<sup>c</sup> Population specific quality control filtering including removing SNP with missing genotypes >0.60, minor allele frequency (MAF) <0.01, Hardy-Weinberg Equilibrium (HWE) P-value < 1e-06, Illumina score <0.8 and at least one Mendelian error in POP B.
<sup>d</sup> Last common quality control filtering retaining markers with unique position in the genome, prioritizing SNP with MAF > 0.05 in the three commercial populations and evenly distributed across the genome
<sup>e</sup> Mitochondria
<sup>f</sup> Unplaced scaffolds

### Animals ethics approval

DNA sampling was carried out in accordance with the commercial practice and norms by Aquacorporación Internacional and Aquamerica.

## RESULTS

### SNP discovery

Whole-genome resequencing of 326 fish yielded a mean of 79.6 (SD = 65.0) millions of raw reads per fish, with a minimum and maximum of 20.6 and 545.6 millions of raw reads per fish, respectively. Quality controlled reads were aligned to the Nile tilapia reference genome, and an average of 76.3 (SD = 64.6) million read per fish, with a minimum and maximum of 20.5 and 543.1 million reads per fish, respectively, could be confidently and uniquely mapped to a single position in the genome and these were used for SNP discovery. Thus, the mean coverage for each fish was 8.7x (SD = 8.9x), with a minimum and maximum of 2.1x and 65.7x coverage per fish, respectively. After the SNP discovery phase, approximately 38,454,404 sequence variants were identified across the panel of 326 individuals. A total 29,956,401 non-redundant SNPs were identified across the panel of 326 fish, and 26,415,097 (88.17%) of these SNPs passed the genotypes quality (GQ<0.15) and minimum quality (minQ<40) filters (Table 1). After discarding 1,596 SNPs from mitochondria, specific QC filters were applied for each population separately, removing SNP based on missing genotypes >0.60, MAF <0.01, HWE P-value < 1e-06, at least one Mendelian error assessed in trios from POP B and non-unique position of SNP probes in the Nile tilapia genome. A total of 261,550; 887,072 and 461,645 SNP were retained after the filtering steps mentioned above for POP A, POP B and POP C, respectively. From all these high quality SNP variants, only 31,694 were common between the three populations and 238,025 SNP variants were common between the two high priority populations POP B and POP C. After applying THIN < 9 kb command in order to select SNP as evenly distributed along the genome as possible only 16,275 SNP were common between the three populations, which were used as the base. The gaps to have a mean of one SNP every 9 kb were filled with additional 33,769 SNP common between POP B and POP C to reach a total of 50,044. Out of these 50,04 SNPs, 44 SNPs from short unplaced scaffolds were removed.

### SNP distribution and annotation

To determine the distribution of SNPs in the Nile tilapia genome, we identified their chromosome and position into the public GenBank accession assembly GCF_001858045.1 produced by the University of Maryland and the University of Stirling (Conte et al., 2017). The SNPs cover 1.01 Gb of the total assembly length and averaged one SNP every 9 kb. A total of 47,349 SNP (94.70%) were located in chromosomes and 2,651 SNP were located into unplaced scaffolds. After SNP annotation, we found that most of the uniquely anchored SNPs were located in introns (57.81%). Further a total of 12.2%, 11.97%, 7.16% and 0.63 were located downstream, upstream, intergenic and exon regions, respectively. The remaining SNPs were found in splice acceptor, splice donor, splice site, 3’UTR and 5’UTR regions. The Pearson correlation coefficient between the number of SNPs within each chromosome and total chromosome size in terms of Mb is *r* = 0.95 (p-value < 2.24e^−11^). The relationship between the number of SNPs per chromosome and the total chromosome length in Mb is shown in Figure 1. Thus, the discovered SNP present an even distribution across the chromosomes on the Nile tilapia genome assembly.

**Fig. 1.**
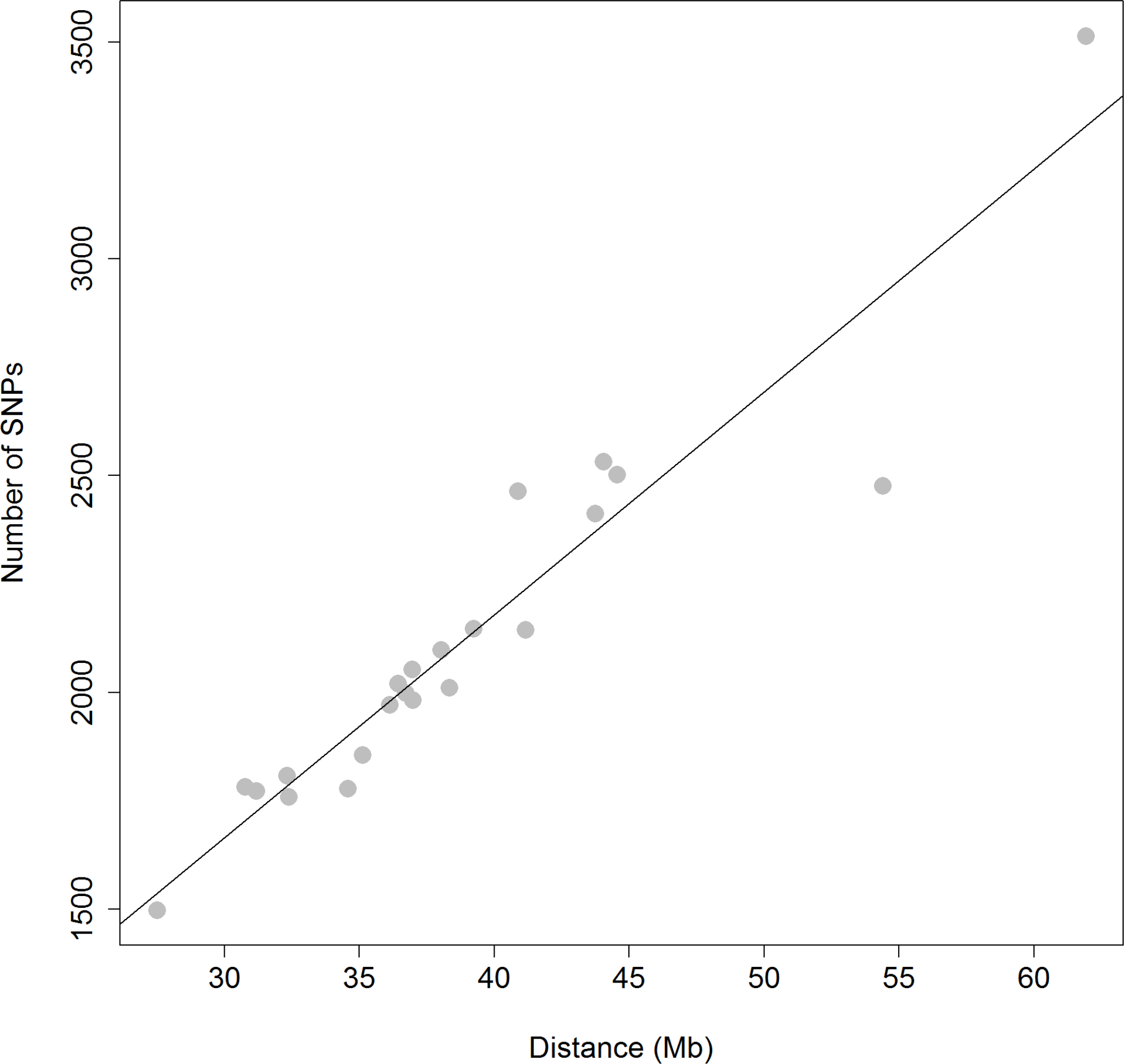
Relationship between the number of SNPs and chromosome length. Scatter plot of the number of SNPs per chromosome and the total chromosome length in Mb according to the assembly GCF_001858045.1. The correlation coefficient between the number of SNPs and chromosome size is *r* = 0.95.

### SNP validation and population segregation

We also performed comparisons between different populations in terms of population genetic estimates using a 50K SNP validation panel. In this respect, the percentage of SNP segregating in HWE in all the populations was 99%, 98% and 94% of the 50K SNP validation panel for POP A, POP B and POP C, respectively. Furthermore, these SNPs showed 76% and 87%, 89% and 93%, and 90% and 93% of MAF > 0.05 and MAF > 0.01 for POP A, POP B and POP C, respectively (Table 2). The distribution of MAF values across SNPs ranged from 0.04 to 0.50 with mean MAF value of 0.24 ± 0.12 (Figure 2). The average observed and estimated heterozygosity (H_O_ and H_E_) was evaluated in each population (Table 2). Although the H_O_ values were very similar among populations, POP A and POP B expressed the lowest (0.20) and the highest (0.25) H_O_ values, respectively, suggesting that these populations are the least and the most genetically diverse populations in the present study. In the three populations, H_O_ diverged considerably from H_E_, resulting in a heterozygote deficiency compared to HWE expectations.

**Table 2.** Descriptive results of population genetic estimates and statistics for the different populations of farmed Nile tilapia using the 50 K SNP validation panel.

| Population | HWE<sup>a</sup> |  | MAF > 0.05<sup>b</sup> |  | MAF > 0.01<sup>c</sup> |  | H<sub>O</sub><sup>d</sup> | H<sub>E</sub><sup>e</sup> |
| --- | --- | --- | --- | --- | --- | --- | --- | --- |
|  | n | % | n | % | n | % |  |  |
| POP A | 46,139 | 99.07 | 37,843 | 75.69 | 43,450 | 86.9 | 0.2011 | 0.2843 |
| POP B | 45,757 | 98.25 | 44,696 | 89.39 | 46,570 | 93.14 | 0.2497 | 0.3130 |
| POP C | 43,869 | 94.20 | 45,171 | 90.34 | 46,570 | 93.14 | 0.2463 | 0.3243 |

<sup>a</sup>SNPs in Hardy-Weinberg Equilibrium
<sup>b</sup>SNPs with Minor Allele Frequency > 0.05
<sup>c</sup>SNPs with Minor Allele Frequency > 0.01
<sup>d</sup>Observed heterozygosity
<sup>e</sup>Expected heterozygosity

**Fig. 2.**
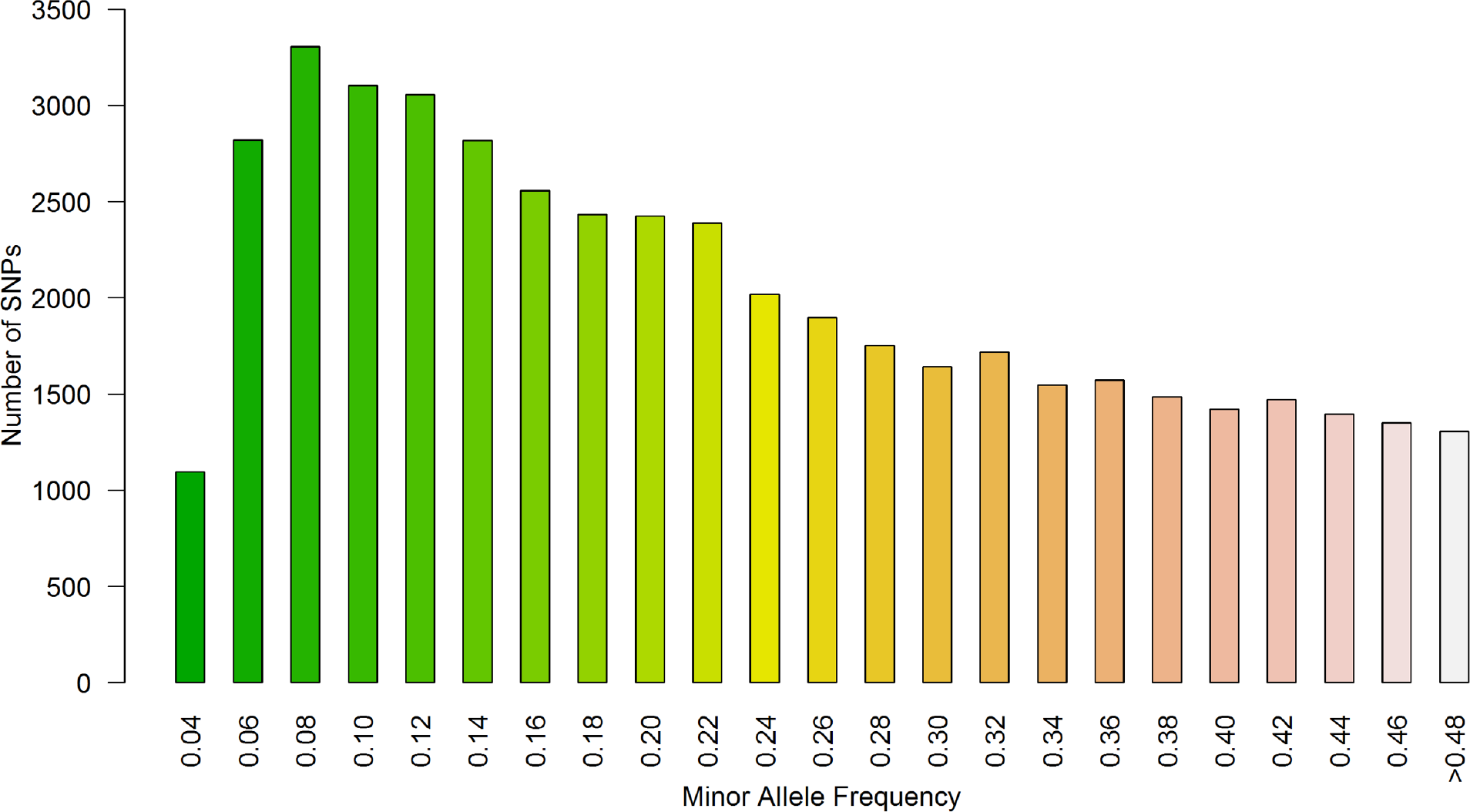
Distribution of minor allele frequencies (MAFs). Distribution of MAFs for the 50K SNP validation panel from 326 samples.

## DISCUSSION

The application of molecular markers into breeding programs has been widely spread along terrestrial and aquaculture species. Dense SNP panels have been shown to facilitate genome-scale studies by allowing the simultaneous evaluation of thousands of SNPs in commercially important fish species, such as Atlantic salmon (Houston et al. 2014; Yáñez et al. 2016) and rainbow trout (Palti et al. 2015). These markers have facilitated the discovery of genetic variants associated with important commercial traits and also the evaluation and implementation of genomic selection in aquaculture species (Correa et al. 2015; Palaiokostas et al. 2016; Bangera et al. 2017; Gutierrez et al. 2018; Vallejo et al. 2018). However, and despite Nile tilapia is widely produced in several countries, with the existence of more than 20 breeding programs (Neira 2010), there are still scarce studies aiming at the application of genome-wide SNP information for the identification of quantitative trait loci and the evaluation and practical implementation of genomic predictions in this species. The SNP discovery strategy used here allowed us to identify a large number of high quality SNPs that can reliably be genotyped across different populations of farmed Nile tilapia with a GIFT origin. The GIFT strain is the most spread Nile tilapia strain used for farming purposes worldwide (Ponzoni et al. 2011). The results from the segregation of SNPs between different populations indicate that the molecular markers identified in the present study would be useful for genetic studies across populations, although the performance of this set of markers would slightly decrease when used in POP A. This is most likely due to the genetic differentiation between populations associated with their distinct origin (founder effect) and independent genetic selection by more than ten generations. The emphasis placed on including SNPs segregating in POP B and POP C may have caused ascertainment bias, which most likely contributed to the lower diversity observed in the POP A. In addition, there is difference in the number of SNPs with MAFs higher than 0.05 and 0.01 for POP A compared to POP B and C. Therefore, these considerations must be taken into account when using the current SNP panel in farmed Nile tilapia populations with different origins and even on wild populations.

A recent study has shown the development of a 58K SNP array for Nile tilapia by means of SNP discovery performed using whole-genome resequencing data of 32 fish from one commercial population (Joshi *et al*., 2018). In this previous study, 40,549 (69.35%) out of 58,466 SNPs were retained after filtering by MAF ≤ 0.05. In our study, between 37,843 (75.68%) and 45,171 (90.34) out of the 50K SNP validation panel were retained after filtering by HWE and MAF ≤ 0.05, indicating a better proportion of SNP validated and a moderate variation (~15%) of availability of SNPs, depending on the target population. The latter is most likely due to ascertainment bias in SNP discovery and selection and it has to be taken into account in further applications of this SNP panel in populations with different origins. When comparing the SNP list from the 50K SNP validation panel against the 58K SNP array developed by Joshi *et al*. (2018), by means of aligning SNP probes, we found that 100% of the SNPs were exclusive to each SNP panel. The high proportion of SNPs exclusive to each of the two SNP panels can be mainly explained by the different genetic background of populations and design of the whole-genome resequencing experiments used for SNP discovery. The 50K SNP validation panel presented here was produced using whole-genome resequencing of 326 fish from three independent populations, which allowed us to have an initial list of 29.9 million putative SNPs, which was almost a three times larger initial set when compared against the previous study from Joshi et al. (2018), in which 32 fish from a single population were whole-genome resequenced, generating 10.5 million putative SNPs for further filtering steps. More importantly, the results presented here indicate that currently available Nile tilapia SNP panels can be considered more as being highly complementary than redundant in terms of the variants represented.

The SNP panel presented here provides an excellent resource for the development of genome-scale studies of biologically and economically important traits. For instance, a recent genome-wide association study using a subset 2.4 million SNPs derived from the 29.9 million SNPs available from the present study, confirmed the anti-Müllerian hormone as a major gene associated with sex determination in different populations of farmed Nile tilapia (Caceres et al. 2019). This information could assist future strategies aiming at generating monosex (all-male) Nile tilapia populations for farming purposes without using hormones, to better exploit the sexual dimorphism present in the species, in which male individuals growth faster than females (Baroiller and D’Cotta 2001; Alcantar et al. 2014). In addition, the SNP panel developed in the present study will also allow the practical implementation of genomic predictions in Nile tilapia selective breeding programs, as it has been reported in a recent study in which an increase in accuracy of EBVs has been demonstrated through the incorporation of genomic information into genetic evaluations for fillet traits (Yoshida et al. 2019a). Finally, the SNP resources presented here will also allow other kind of population genetic studies in farmed populations of Nile tilapia using dense genome-wide information, as for example, has been recently done by the determination of the genetic structure and linkage disequilibrium in farmed populations using dense SNP genotypes (Yoshida et al. 2019b).

## CONCLUSIONS

This paper describes the simultaneous discovery and validation of SNP markers in Nile tilapia through the use of whole-genome resequencing of hundreds of animals. The SNPs identified here will provide an opportunity for the dissection traits of biological and economic importance, such as growth, carcass quality and disease resistance traits, through the application in genome-scale studies. Furthermore, it will allow increasing the response to selection for these traits by means of genomic selection in breeding programs. We believe that downstream applications of this important genomic platform will help to enhance Nile tilapia production by making it more efficient and sustainable.

## Acknowledgments

This study was partially funded through financial support from CORFO grant number 14EIAT-28667 from the Government of Chile. This work was supported by Basal grant of the Center for Mathematical Modeling AFB170001 (UMI2807 UCHILE-CNRS) and Center For Genome Regulation Fondap Grant 15090007. Powered@NLHPC This research was partially supported by the supercomputing infrastructure of the NLHPC (ECM-02). We would like to acknowledge to AquaAmerica and Aquacorporación Internacional for kindly providing the samples used in this work. We would also like to acknowledge to Gabriel Rizzato and Natalí Kunita from AquaAmerica and Diego Salas and José Soto from Aquacorporación International and for their contribution of samples from Brazil and Costa Rica, respectively.

## Author Contributions

J.M.Y. conceived and designed the study, contributed to the analysis and drafted the manuscript. G.Y. contributed to analysis and writing. A.B. drafted the first version of the manuscript. G.C., M.E.L. and A.J. participated in data collection, purification and management of the samples for sequencing and genotyping. R.P., D.D., D.T., and A.M. performed the bioinformatics analysis and contributed to writing. J.P.L. participated in the design of the study and writing. JS and DS contributed to samples collection. All authors have reviewed and approved the manuscript.

## Conflict of Interests

Two commercial organizations (Aquainnovo and Illumina) were involved in the SNP identification and preparation of the manuscript. However, this does not alter public accessibility to data from the SNP data presented in this study. JPL was employed by Benchmark Genetics Chile during the course of the study.

## Data Availability

The sequence data used for SNP discovery will be deposited in public database upon acceptance. The full SNP list can be found in the Figshare repository (accession number 10.6084/m9.figshare.7581581).

